# Identifying neurophysiological features associated with anesthetic state in newborn mice and humans

**DOI:** 10.1101/451831

**Authors:** Mattia Chini, Sabine Gretenkord, Johanna K. Kostka, Jastyn A. Pöpplau, Laura Cornelissen, Charles B. Berde, Ileana L. Hanganu-Opatz, Sebastian H. Bitzenhofer

**Affiliations:** Developmental Neurophysiology, Institute of Neuroanatomy, University Medical Center Hamburg-Eppendorf, Hamburg, Germany; Department of Anesthesiology, Critical Care and Pain Medicine, Boston Children’s Hospital, Boston, Massachusetts; Department of Anaesthesia, Harvard Medical School, Boston, Massachusetts

## Abstract

**One Sentence Summary:** Machine learning reveals consistent features of anesthetic states assessed by intracranial recordings in newborn mice and multichannel EEG in human neonates and infants.

**Abstract:** Monitoring the hypnotic component of anesthesia during surgeries is critical to prevent intraoperative awareness and reduce adverse side effects. For this purpose, electroencephalographic methods complementing measures of autonomic functions and behavioral responses are in use in clinical practice. However, in human neonates and infants existing methods may be unreliable and the correlation between brain activity and anesthetic depth is still poorly understood. Here, we characterize the effects of different anesthetics on activity of several brain areas in neonatal mice and develop machine learning approaches to identify electrophysiological features predicting inspired or end-tidal anesthetic concentration as a proxy for anesthetic depth. We show that similar features from electroencephalographic recordings can be applied to predict anesthetic concentration in neonatal mice, and human neonates and infants. These results might support a novel strategy to monitor anesthetic depth in human newborns.

## Introduction

Reliable monitoring of anesthesia depth is critical during surgery. It allows for loss of consciousness, analgesia and immobility without incurring the risk of side effects and complications due to anesthetic misdosing. Typically used measures to monitor anesthesia depth are inspired and end-tidal anesthetic concentrations as well as physiologic parameters, including respiratory rate and depth (in the absence of neuromuscular blockade or controlled ventilation), heart rate, blood pressure, and responses to noxious stimuli (*1*). These measures all respond to spinal and brainstem reflexes and are not specific for arousal or cortical responses to noxious events.

Anesthesia-induced changes in brain activity can be measured with electroencephalographic (EEG) recordings. Specific algorithms have been developed to predict anesthesia depth in adults (*2-4*). The most commonly used of such methods, the Bispectral Index, has been shown to significantly reduce intraoperative awareness, amount of anesthetic used, recovery time and post-anesthesia care unit stay in a recent Cochrane meta-analysis (*5*), but see (*6, 7*). However, evidence of similar benefits in infants and younger children is sparse, as recently shown (*8-10*). EEG in anesthetized infants changes dramatically depending on postnatal age (*8, 11-14*).

EEG recordings mainly monitor neocortical activity. Converging evidence from animal and human studies has shown that most anesthetics slow electroencephalographic oscillations (*15-17*). While power at high frequency oscillations is reduced (>40 Hz), power at slower frequencies (<15 Hz) is enhanced (*15*). The computations underling proprietary indexes such as the Bispectral index or Narcotrend are thought to take advantage of these phenomena (*18*). However, in preterm and term neonates for the first weeks of life, EEG during sleep-wake cycles is weakly correlated with behavioral states and shows characteristic bursts or spontaneous activity transients (*19, 20*). Anesthesia-induced theta and alpha oscillations have been reported to emerge around 3-4 months of age, albeit with less frontal predominance than in older children and adults (*8, 10*). Moreover, high concentrations/doses of anesthetics have been reported to depress brain activity and enhance signal discontinuity in both human and rodent neonates (*9, 21,* 22). However, to our knowledge, a comprehensive algorithmic approach identifying electroencephalographic parameters that robustly correlate with anesthetic depth during early postnatal development is still lacking.

Here, we developed a novel strategy to model anesthesia depth by using common electrophysiological features that correlate with inhaled anesthetic concentrations during early development in age-matched mice and humans. We performed intracranial electrophysiological recordings to study the temporal and dose-dependent dynamics of brain activity in neonatal mice (postnatal day (P) 8-10) during bolus urethane administration, and during dose-titrated isoflurane general anesthesia, respectively. Dominant local field potential (LFP) features of anesthetic state were identified and used to develop a machine-learning algorithm that distinguishes non-anesthetized from deeply anesthetized states, and predicts anesthetic concentration as a proxy for anesthetic depth. Using a similar approach, we used multielectrode EEG recordings to study the dose-dependent dynamics of brain activity in a secondary analysis of a combined new and previously reported data set (*10*) of human infants 0-6 months of age during induction, maintenance and emergence from general anesthesia (sevoflurane, isoflurane, or desflurane) administered for routine surgical care. Dominant EEG features of anesthetic state were identified and used to develop a machine-learning algorithm to predict end-tidal volume anesthetic concentration (an indirect measure of anesthetic concentration in the brain, and anesthetic depth).

## Results

### Anesthesia affects the occurrence but not the spectral and temporal structure of oscillatory events in neonatal mice

We monitored the impact of anesthesia on immature brain activity in several cortical areas (prefrontal cortex (PFC), hippocampus (HP), and lateral entorhinal cortex (LEC)) as well as in a sensory area (olfactory bulb (OB)). For this, multi-site extracellular recordings of LFP and multi-unit activity (MUA) were performed from P8-10 mice before and for 45 minutes after induction of anesthesia by intraperitoneal urethane injection (Fig. 1A), an anesthetic commonly used in rodents (*23, 24*).

**Fig. 1.**
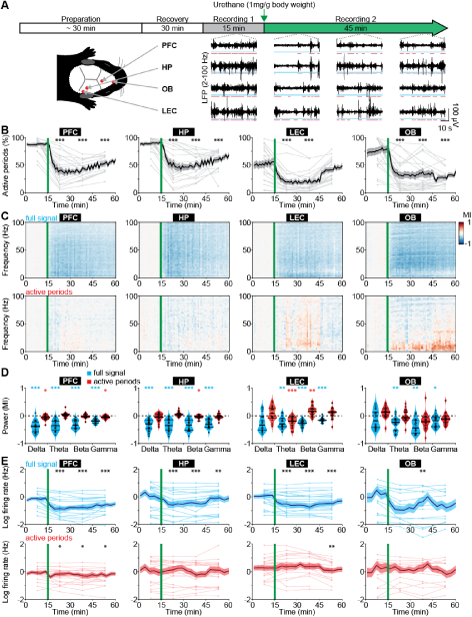
Frequency-unspecific dampening of neuronal activity during urethane anesthesia in neonatal mice. (**A**) Schematic representation of experimental paradigm and recording sites as well as characteristic LFP recordings of discontinuous activity in the PFC, HP, LEC, and OB of neonatal mice (P8–10) during non-anesthetized and urethane-anesthetized state. Time windows of active periods are marked by red lines. (**B**) Line plots displaying the relative occurrence of active periods normalized to total recording time in PFC, HP, OB and LEC before and after urethane injection. (**C**) Color-coded MI of power spectra for full signal (top) and active periods (bottom) recorded in PFC, HP, LEC and OB of neonatal mice before and after urethane injection. (**D**) Violin plots displaying the MI of power in delta (2-4 Hz), theta-alpha (4-12 Hz), beta (12-30 Hz) and gamma (30-100 Hz) frequency bands for full signal (blue) and active periods (red) recorded in the PFC, HP, LEC and OB. (**E**) Line plots displaying MUA rates during full signal (blue) and active periods (red). In (B), (C) and (E) green lines correspond to the time point of urethane injection.

The recorded network activity had a highly fragmented structure (defined as discontinuous activity) in all investigated areas (PFC, HP, LEC and OB). The full signal (i.e. entire LFP trace) consisted of transient episodes of oscillatory discharges with mixed frequencies (from here referred to as ‘active periods’), alternating with periods of relative electrical silence and suppressed activity (from here referred to as ‘silent periods’) (Fig. 1A) (*23, 25-28*). The prevalence of active periods decreased rapidly and robustly over time in all investigated brain areas upon urethane injection (Fig. 1B). The most prominent reduction was observed 5 to 15 minutes after urethane injection. A partial recovery towards baseline levels during the following 30 minutes was detected in cortical areas, and to a lesser extent in OB (Fig. 1B). The temporal sequence of events likely reflects the pharmacokinetics of urethane and is line with the previously reported long-lasting effects of urethane anesthesia (*29*).

The anesthesia-induced reduced occurrence of active periods was reflected in a broadband (1-100 Hz) decrease in oscillatory power shown as modulation index (MI) defined as (power_post_-power_pre_) / (power_post_+power_pre_). In contrast, power spectra during active periods were largely unaffected (Fig. 1C). Spectral properties of full signal and active periods were quantified for delta (2-4Hz), theta-alpha (4-12 Hz), beta (12-30 Hz) and gamma (30-100 Hz) frequency bands for the first 15 minutes post urethane administration. In contrast to the significant reduction of full signal power in all frequency bands, the power during active periods was only marginally affected by anesthesia (Fig. 1D). Thus, urethane anesthesia affected network activity in the immature rodent brain predominantly by decreasing the amount of active periods without perturbing the frequency structure of active periods. This is in stark contrast with the well-characterized switch from a low-amplitude high-frequency regime to a high-amplitude low-frequency regime of electrical activity that has been reported for the adult rodent and human brain (*17, 30*).

Anesthesia was shown to induce alterations of long-range network interactions in adult rodents (31) and humans (*32-34*). We examined whether similar alterations are present in the immature mouse brain. Simultaneous recordings of HP and PFC, as well as OB and LEC were analyzed to assess the effects of anesthesia on long-range functional coupling. We previously showed that at the end of the first postnatal week hippocampal theta bursts drive the oscillatory entrainment of local circuits in the PFC, whereas discontinuous activity in OB controls the network activity in LEC (*26, 27, 35*). Urethane did not modify these interactions. The synchrony within networks quantified by HP-PFC and OB-LEC coherence was similar during baseline (no urethane anesthesia) and in the presence of urethane (Fig. S1A). These data indicate that the core features of long-range functional coupling are retained under anesthesia in neonatal mice.

Anesthesia modified neuronal firing in all investigated areas. Firing rates in PFC, HP, LEC and OB decreased after urethane injection and only partially recovered during the following 45 min (Fig. 1E). However, firing rates during active periods were only marginally affected. To examine whether the timing of neuronal firing to the phase of oscillatory activity was altered by anesthesia, we calculated pairwise phase consistency (PPC), a firing rate-independent measure of spike-LFP phase locking (*36*). All four brain regions showed similar frequency-resolved phase locking profiles before and after urethane injection (Fig. S1B,C).

Anesthetics have been shown to alter the excitation/inhibition balance in the adult brain through their action on specific ion channels involved in synaptic transmission (*37*). Such alteration is usually monitored by changes in the 1/f slope of power spectral density. Further, signal complexity and information content measured by sample entropy have been correlated with behavioral states of adults, such as consciousness, sleep/wake states and anesthesia (*38, 39*). For neonatal mice, we observed similar values of 1/f slope and sample entropy before and during urethane anesthesia (Fig. S1D-F), suggesting that urethane does not perturb cortical excitation/inhibition balance and signal complexity at this early age. The findings provide additional evidence to the hypothesis that anesthesia has unique effects on the immature brain.

To add additional evidence for this hypothesis, we extended the time window of investigation and performed extracellular recordings from the PFC of juvenile mice (P24-39). In contrast to the frequency-unspecific reduction of active periods in neonates, urethane anesthesia increased the oscillatory power in the delta frequency band and suppressed power in beta and gamma frequency bands (Fig. S2), confirming the anesthetic effects in the adult brain (*15-17*).

Taken together, these results indicate that urethane anesthesia dampened neonatal brain activity mainly by augmenting the discontinuity of network activity, i.e. reducing the proportion of time the brain spent in active periods. However, the active periods were largely unaffected in their temporal structure and firing dynamics. In contrast, urethane anesthesia in older mice led to frequency-specific changes. Thus, urethane anesthesia differently impacts neonatal and adult brain activity in mice.

### Suppression of active periods predicts anesthetic concentration in neonatal mice

To test whether the effects of urethane on neonatal brain activity generalize to other anesthetics, we performed LFP and MUA recordings from HP and PFC of P8-10 mice at increasing doses of isoflurane-induced anesthesia (0, 1, 2 and 3%; 15 min per concentration) (Fig. 2A). Isoflurane reduced the incidence of active periods in a dose-dependent manner (Fig. 2B). Accordingly, the broadband reduction of LFP power was also dependent on isoflurane concentration (Fig. 2C,D). Power spectra of active periods remained largely unaffected in the presence of isoflurane, similarly to the urethane effects (Fig. 2C,D). MUA rates during active periods in PFC and HP were hardly modified in the presence of isoflurane, yet the overall firing decreased corresponding to the reduced occurrence of active periods (Fig. 2E). Together, these findings identify the suppression of active periods as the main effect of bolus urethane injection and isoflurane anesthesia in the neonatal mouse brain.

**Fig. 2.**
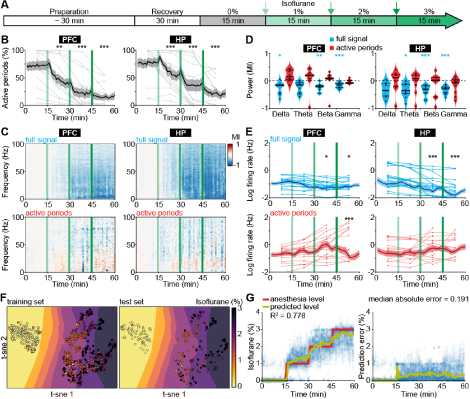
Suppression of active periods in relationship with the depth of isoflurane anesthesia in neonatal mice. (**A**) Schematic representation of experimental protocol for LFP recordings without anesthesia and during increasing levels of isoflurane anesthesia in neonatal mice (P8-10). (**B**) Line plots displaying the relative occurrence of active periods in PFC and HP during increasing levels of isoflurane anesthesia. (**C**) Color-coded MI of power spectra for full signal (top) and active periods (bottom) during increasing levels of isoflurane anesthesia. (**D**) Violin plots displaying the MI of power in delta (2-4 Hz), theta (4-12 Hz), beta (12-30 Hz) and gamma (30-100 Hz) frequency bands for full signal (blue) and active periods (red). (**E**) Line plots displaying MUA firing rates during full signal (blue) and active periods (red). In (B), (C) and (E) green lines correspond to the time points of increasing isoflurane anesthesia. (**F**) Visualization of anesthesia depth prediction by t-sne plots. Background color codes for predicted anesthesia depth, while the color of the dots represents the actual anesthesia level in the training (left) and test set (right). (**G**) Scatter plots displaying anesthesia depth predictions with support vector regression (left) and absolute errors between anesthesia depth prediction and actual anesthesia depth (right).

The development-specific response of the immature brain to anesthesia might represent the main obstacle when trying to predict anesthesia depth in infants using algorithms based on the mature brain activity of adults. Therefore, we next aimed to use electrophysiological properties specific for anesthetized neonatal mice to predict the concentration of administered isoflurane. We used support vector regression (Fig. S3), with the following input features: median amplitude of broadband LFP, percent of time spent in active periods, and spectral power from 1 to 100 Hz in 10 Hz bins for both hippocampal and prefrontal activity. An additional feature was the output of a support vector classifier that received the same features as for the support vector regression, and that was designed to predict whether the animal was under anesthesia or not. The algorithm accurately predicted anesthesia depth across all levels of isoflurane concentration (Fig. 2F,G). Estimation of information content of the different features identified the median amplitude of broadband LFP as the most informative feature (Fig. S4A). As the power of active periods was only marginally affected by anesthesia, this feature mainly mirrors the suppression of active periods. Interestingly, the algorithm was also able to distinguish non-anesthetized from anesthetized recordings from neonatal mice under urethane, even though it had not been exposed to this dataset during training (Fig. S4B).

Thus, features of electrophysiological activity that capture the particularities of immature neuronal networks can predict anesthetic concentration in neonatal mice. The generalization of the classifier to a different anesthetic indicates that it can identify general anesthesia-related features of brain activity in neonatal mice.

### Frequency-unspecific suppression of activity in anesthetized human neonates and young infants

To test if human neonates and infants, similarly to mice, respond to anesthesia with a broadband decrease of periods of oscillatory activity, we examined EEG recordings from humans aged 0-6 months postnatal age, who received general anesthesia with volatile anesthetics (sevoflurane 32 subjects, isoflurane 2 subjects, desflurane 1 subject) for surgery (Tab. S1).

In neonatal mice, the median LFP amplitude of broadband activity was identified as the most informative feature to predict anesthetic depth. We therefore applied the same data analysis approach to human EEG data (Fig. S5). We found the median amplitude of broadband EEG activity (averaged across all recording electrodes across the scalp) was negatively correlated with endtidal anesthetic concentration (etAnesthetic) in human neonates from birth until 2 months postnatal age (Fig. 3A,B). For older human infants, the correlation of the median EEG amplitude with the anesthetic concentration switched to a positive correlation, in agreement with adult human data (*40*). This relationship was even stronger using expected birth age, corrected for conceptional age (Fig. S6A). This switch from negative to positive correlation was also visible in the normalized median EEG amplitude when averaged for age-grouped babies (0-2, 2-4, 4-6 months) (Fig. 3C).

**Fig. 3.**
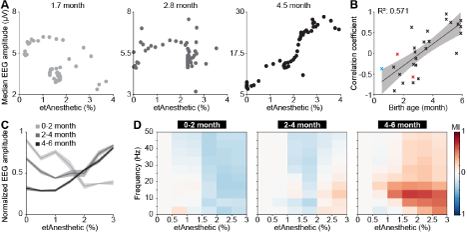
Age-dependent switch from broadband suppression to frequency-specific effects of general anesthesia on EEG activity in human neonates and infants. (**A**) Scatter plots displaying the median EEG amplitude as a function of anesthetic concentration for representative examples of 0-2, 2-4 and 4-6 months of age. (**B**) Scatter plot displaying the correlation coefficient of median EEG amplitude and anesthetic concentration in relationship to birth age for sevoflurane (black), isoflurane (red), and desflurane (blue). (**C**) Line plots displaying normalized EEG amplitude as a function of anesthetic concentration. (**D**) Color-coded MI of median EEG amplitudes in different frequency bands as a function of anesthetic concentration for human babies of 0-2 months (left), 2-4 months (middle) and 4-6 months (right).

Quantification of median EEG amplitude across frequencies revealed a broadband suppression of EEG activity in human neonates of 0-2 months (Fig. 3D). In contrast, the relationship between activity amplitude and etAnesthetic indicated frequency-specificity in human infants of 2-4 and 4-6 months, as previously reported (*9*). Frontal activity has been shown to be particularly sensitive to age-varying anesthesia-related effects in human neonates (*8*). Analysis of only frontal electrodes (Fp1, Fp2, F3, F4, F7, F8, Fpz) showed the same age-dependent anesthesia-induced changes as analysis of full scalp electrodes (Fig. S6B-D).

Thus, analogous to what we found in neonatal mice, general anesthesia in human infants younger than 2 months suppressed neuronal population activity, as reported previously (*8*), while at older age anesthesia induced frequency-specific effects.

### A model to predict end-tidal volume of sevoflurane anesthesia in human neonates and infants

The correlation of EEG activity with etAnesthetic as well as the similar effects of anesthesia in neonatal mice and in humans from birth to 2 months old, suggests that anesthetic depth in babies might be predicted using similar features to those used in neonatal mice. To test this, we used a machine-learning algorithm with a similar architecture as the one we developed for neonatal mice (Fig. S3). The algorithm was modified to account for the developmental switch from broadband suppression to frequency-specific modulation by training three different regressors using 2 and 4 months as cut-offs. All regressors received the same input features (see Methods and Fig. S5). Features derived from EEG activity were able to predict etAnesthetic with high accuracy for all age groups (0-2 months R^2^=0.806, 2-4 months R^2^=0.688, 4-6 months R^2^=0.787) (Fig. 4A-C). In line with the frequency-specific alterations observed only in the older age groups, frequency-related features were rated more important for prediction of anesthesia depth in infants of 2-4 and 4-6 months than in neonates of 0-2 months (Fig. S7A-C). Predicting anesthesia depth for all ages with a single classifier considering age as an input feature performed with high accuracy (0-6 months R2=0.689) (Fig. 4D, S7D). This result confirms the age-varying effects of anesthesia on the brain and stresses the importance of considering age when developing algorithms aiming to assess anesthetic depth.

**Fig. 4.**
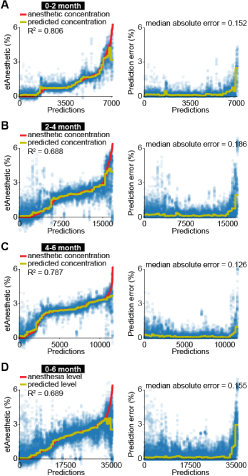
EEG activity is predictive for anesthetic concentration in human infants. (**A**) Scatter plots displaying anesthetic concentration predictions of support vector regression (left) and absolute errors between anesthetic concentration prediction and actual anesthetic concentration (right) for human neonates of 0-2 months. (**B**) Same as (A) for human infants of 2-4 months. (**C**) Same as (A) for human infants of 4-6 months. (**D**) Same as (A) for human neonates and infants of 0-6 months.

Thus, mouse and human neonates show similar changes in network activity in response to anesthesia. These results highlight how neurophysiological activity could be beneficial for future attempts at predicting anesthetic depth in clinical settings.

## Discussion

Monitoring brain function during anesthesia is desirable to avoid intraoperative awareness and side effects resulting from unnecessarily high doses of anesthetics. Since consciousness is an elusive concept and cannot be directly measured, EEG features have been used to guide anesthesia delivery during human surgery. Monitoring methods developed for adults perform poorly in human neonates and infants, particularly during the first months of life (*11-13, 41*). Age-specific effects of anesthetics on immature brain activity are considered the main reason for such poor performance. Implementation of neonate- and infant-specific anesthesia monitors requires elucidation of distinct anesthesia-induced EEG features during early development. We took advantage of a translational approach to address this open question. We first carried out an in depth investigation of anesthesia effects on brain activity in neonatal mice, and then applied this knowledge to develop features that would correlate with anesthetic concentration in human neonates.

In contrast to the continuous EEG signal observed in adults, neonatal EEG around birth is characterized by a highly discontinuous and fragmented temporal organization, with bursts of cerebral activity (active periods) alternating with interburst intervals lacking activity (silent periods) (*42-48*). Neonatal mice show a similar discontinuous organization of cortical activity (*23, 25, 26*). In accordance with the similar organization of early activity patterns in age-matched mouse pups and human infants, we found comparable effects of anesthesia on LFP and EEG signals, respectively.

It is well established that in the adult rodent and human brain most anesthetics favor slow oscillations at the expense of faster ones, thereby slowing the electroencephalographic rhythm (*15-17*). This principle is thought to underlie most algorithms that are clinically used to predict anesthesia depth (*13*). Indeed, such algorithms perform poorly with anesthetics, such as ketamine, that do not share this mechanism of action (*49*). In line with previous studies (*50, 51*), we report that both urethane and isoflurane anesthesia affect brain activity in a different way in neonatal mice. Instead of favoring slow oscillations at the expense of faster ones, anesthesia in neonatal mice broadly suppresses activity in a frequency-unspecific manner. The dampening of cortical activity for human infants of 0-2 months suggests a development specific effect of anesthesia on immature brain activity that translates between mice and humans.

In rodents, the switch from activity suppression to frequency-specific modulation of neuronal activity by anesthesia has been reported to occur around P12 (*50*). This coincides with the emergence of slow oscillations during sleep, suggested to depend on the maturation of thalamocortical networks (*50, 52*). Consistent with our previous studies evaluating EEG properties of this data set, we found that theta and alpha oscillatory activity under anesthesia emerges in humans at around 4 months postnatal age (*8-10*). Future studies with an increased age range in mice and humans, including data of human infants studied at preterm, and children in older than 6 months of age, may deepen the understanding of anesthetic effects on brain activity throughout development.

The anesthetics evaluated across species in this study were comparable but not identical in terms of mechanism of action. Moreover, anesthetic management practices used in mice were simplified compared to commonly-used anesthetic practices in the clinic. Multimodal anesthesia requires the use of low-dose anesthetics in combination with analgesic and neuromuscular blocking agents to provide optimal anesthesia and reduce side effect. These agents act on different drug targets in the nervous system and may have subtle but different effects on brain oscillatory activity (*53*).

In adult human volunteers, the correlation with anesthetic depth and EEG parameters can be performed using verbal reports to establish a threshold for unconsciousness (*15*). However, in non-verbal populations such as human infants, one must rely on indirect behavioral measures which are more readily performed on emergence rather than induction and incision (*54*). Future investigations need to include surgical incision and other stimuli into the mouse models to understand with greater granularity the anesthetic titration around the minimal concentrations required to suppress movement, autonomic, and cortical responses to noxious stimuli.

In summary, we report that the suppression of brain activity in mouse and human neonates correlates with anesthetic concentration. The detailed understanding of anesthesia effects on network activity in mice allowed us to identify features and develop a machine-learning algorithm that is able to predict anesthetic concentration from EEG recordings in human neonates. We propose that, after appropriate training, an algorithm based on what we introduce here could learn to associate specific EEG effects with certain anesthetic doses. Eventual mismatches between administered and predicted anesthetic dose would then identify patients that are particularly sensitive/insensitive to an anesthetic, thus helping the anesthetist in administering appropriate levels of anesthetics. By these means, the risk of adverse neurodevelopmental outcome might be mitigated.

## Materials and methods

### Animals

All experiments were performed in compliance with the German laws and the guidelines of the European Community for the use of animals in research and were approved by the local ethical committee (G132/12, G17/015, N18/015). Experiments were carried out on C57Bl/6J mice of both sexes. Timed-pregnant mice from the animal facility of the University Medical Center Hamburg-Eppendorf were housed individually at a 12 h light/12 h dark cycle, with ad libitum access to water and food. Day of birth was considered P0.

### In vivo electrophysiology in neonatal mice

Multisite extracellular recordings were performed in the PFC and HP, or LEC and OB of P8-10 mice. Pups were on a heating blanket during the entire procedure. Under isoflurane anesthesia (induction: 5%; maintenance: 2.5%), craniotomies were performed above PFC (0.5 mm anterior to bregma, 0.1-0.5 mm right to bregma) and HP (3.5 mm posterior to bregma, 3.5 mm right to bregma), or LEC (0 mm anterior to lambda, 6.5 mm right to lambda) and OB (0.5-0.8 mm anterior from the frontonasal suture, 0.5 mm right from internasal suture). Pups were head-fixed into a stereotaxic apparatus using two plastic bars mounted on the nasal and occipital bones with dental cement. Multisite electrodes (NeuroNexus, MI, USA) were inserted into PFC (four-shank, A4×4 recording sites, 100 μm spacing, 2.0 mm deep) and HP (one-shank, A1×16 recording sites, 50 μm spacing, 1.6 mm deep, 20° angle from the vertical plane), or LEC (one-shank, A1×16 recording sites, 100 μm spacing, 2 mm deep, 10° angle from the vertical plane) and OB (one-shank, A1×16 recording sites, 50 μm spacing, 1.4-1.8 mm deep). A silver wire was inserted into the cerebellum and served as ground and reference electrode. Pups were allowed to recover for 30 min prior to recordings. Extracellular signals were band-pass filtered (0.1-9,000 Hz) and digitized (32 kHz) with a multichannel extracellular amplifier (Digital Lynx SX; Neuralynx, Bozeman, MO, USA).

### In vivo electrophysiology in juvenile mice

Multisite extracellular recordings were performed in the PFC of P24–39 mice. Under isoflurane anesthesia (induction: 5%; maintenance: 2.5%), a metal head-post (Luigs and Neumann) was attached to the skull with dental cement and 2-mm craniotomies were performed above PFC (0.5-2.0 mm anterior to bregma, 0.1-0.5 mm right to bregma) and protected by a customized synthetic window. A silver wire was implanted in the cerebellum as ground and reference electrode. Surgery was performed at least five days before recordings. After recovery mice were trained to run on a custom-made spinning-disc. For recordings craniotomies were uncovered and multisite electrodes (NeuroNexus, MI, USA) were inserted into PFC (one-shank, A1×16 recording sites, 50 μm spacing, 2.0 mm deep). Extracellular signals were band-pass filtered (0.1-9,000 Hz) and digitized (32 kHz) with a multichannel extracellular amplifier (Digital Lynx SX; Neuralynx, Bozeman, MO, USA).

### Recordings under urethane

Activity was recorded for 15 min without anesthesia before intraperitoneally injecting urethane (1 mg/g body weight; Sigma-Aldrich, MO, USA). Activity was recorded for further 45 min. Animals were transcardially perfused after recordings, brains were sectioned coronally, and wide field images were acquired to verify recording electrode positions.

### Recordings under isoflurane

Mouth piece of an isoflurane evaporator (Harvard apparatus, MA, USA) was placed in front of the pups in the recording setup until animals accustomed to it. Activity was recorded for 15 min 0% isoflurane, but with the evaporator running (1.4 l/min). Afterwards, isoflurane was added to the airflow and increased every 15 min (1, 2, 3 %). Animals were transcardially perfused after recordings, brains were sectioned coronally, and wide field images were acquired to verify recording electrode positions.

### Electroencephalographic recordings in human neonates and young infants

Neonates and infants who were scheduled for an elective surgical procedure were recruited from the pre-operative clinic at Boston Children’s Hospital from 12/2012 to 08/2018 (under Institutional Review Board P-3544, with written informed consent obtained from parents/legal guardians). Subjects required surgery below the neck, were clinically stable on the day of study and American Society of Anesthesiologists’ physical status I or II. Exclusion criteria were born with congenital malformations or other genetic conditions thought to influence brain development, diagnosed with a neurological or cardiovascular disorder, or born at <32 weeks post-menstrual age. Datasets from previously published work (n=25) (*10*) and new subjects (n=10) were included in the analysis. Data are presented from 35 subjects aged 0-6 months.

#### Anesthetic management

Each patient received anesthesia induced with sevoflurane (32 subjects), isoflurane (2 subjects) or desflurane (1 subject) alone, or a combination of one of the previous and nitrous oxide. Epochs used for analysis were comprised of sevoflurane, isoflurane or desflurane administration with air and oxygen, titrated to clinical signs; end-tidal anesthetic concentration was adjusted per the anesthetist’s impression of clinical need, not a pre-set end-tidal anesthetic concentration.

#### EEG recording

EEG data were acquired using an EEG cap (WaveGuard EEG cap, Advanced NeuroTechnology, Netherlands). 33- or 41-recording electrodes were positioned per the modified international 10/20 electrode placement system. Reference and ground electrodes were located at Fz and AFz respectively. EEG activity from 0.1-500 Hz was recorded with an Xltek EEG recording system (EMU40EX, Natus Medical Inc., Canada). Signals were digitized at a sampling rate of 1024Hz and a resolution of 16-bit.

#### Clinical data collection

Demographics and clinical information were collected from the electronic medical records and from the in-house Anesthesia Information Management System (AIMS) (Tab. S1). End-tidal sevoflurane, oxygen, and nitrous oxide concentrations were downloaded from the anesthetic monitoring device (Dräger Apollo, Dräger Medical Inc., PA, USA) to a recording computer in real-time using ixTrend software (ixcellence, Germany). Signals were recorded at a 1 Hz sampling rate.

#### Data analysis

In vivo data were analyzed with custom-written algorithms in the Matlab environment. Data were processed as following: band-pass filtered (500–5,000 Hz) to analyze MUA and band-pass filtered (2-100 Hz) using a third-order Butterworth filter before downsampling to analyze LFP. Filtering procedures were performed in a phase preserving manner.

#### Multi-unit activity

MUA was detected as the peak of negative deflections exceeding five times the standard deviation of the filtered signal and having a prominence larger than half the peak itself. Firing rates were computed by dividing the total number of spikes by the duration of the analyzed time window.

#### Detection of oscillatory activity

Discontinuous active periods were detected with a modified version of a previously developed algorithm for unsupervised analysis of neonatal oscillations (*55*). Briefly, deflections of the root mean square of band-pass filtered signals (1–100 Hz) exceeding a variance-depending threshold were considered as network oscillations. The threshold was determined by a Gaussian fit to the values ranging from 0 to the global maximum of the root-mean-square histogram. If two oscillations occurred within 200 ms of each other they were considered as one. Only oscillations lasting >1 s were included, and their occurrence, duration and amplitude were computed.

#### Power spectral density

For power spectral density analysis, 1 s-long windows of full signal or network oscillations were concatenated and the power was calculated using Welch’s method with non-overlapping windows.

#### Imaginary coherence

The imaginary part of coherence, which is insensitive to volume-conduction-based effects (*56*), was calculated by taking the absolute value of the imaginary component of the normalized cross-spectrum:

#### Pairwise phase consistency

Pairwise phase consistency was computed as previously described (*36*). Briefly, the phase in the band of interest was extracted using Hilbert transform and the mean of the cosine of the absolute angular distance among all pairs of phases was calculated.

#### 1/f slope

1/f slope was computed as previously described (*19*). We used robust linear regression (MATLAB function *robustfit)* to find the best fit over 20-40 Hz frequency range of the power spectral density, in one minute bins.

#### Sample entropy

Sample Entropy was computed using the SampEn function (MATLAB File Exchange) in 1.5 seconds windows and in 2 Hz frequency bins. Tolerance was set to 0.2 * std(signal), and tau to 1.

#### EEG data analysis

EEG signal was visually inspected to detect and reject channels with low signal to noise ratio, and re-referenced to a common average reference. The signal was automatically scored in five seconds epochs, and channels in which signal was significantly contaminated by artifacts (patient handling, surgical electrocautery etc.) were discarded. Epochs were rejected if signal was saturated due to electrocautery, signal exceeded 150μV, or the median signal across all EEG channels exceeds 30μV (Fig. S5). Minutes containing more than 10s of contaminated signal were removed from further analysis. On average 14 +/- 9% (median +/- median absolute deviation) of the signal was discarded. To compute EEG amplitude, we smoothed the absolute value of the signal, using a moving average filter with a window of 1024 points (1 second). If more than one volatile anesthetic was used, we retained only epochs in which the main anesthetic was used in isolation. Subjects with epidural anesthesia halfway through the surgery (n=2 subjects), or with less than 20 minutes of artifact-free signal (n=5 subjects) were excluded from further analysis.

#### Feature engineering

Features to predict anesthetic concentration in neonatal mice were calculated in one minute bins. LFP power in the 1-100 Hz range in 10 Hz bins, the percentage of active periods, median length and number of oscillations, median and maximum signal amplitude were computed. All features were computed for both PFC and HP, and were normalized to their median value in the non-anesthetized 15 minutes of recordings. Features to predict anesthetic concentration in human infants were also calculated in one minute bins. The median amplitude of the smoothed EEG signal, and the percentage of the EEG envelope that fell into each amplitude quartile was computed. Amplitude quartiles were computed on the entire EEG trace, averaged over channels. All features were calculated for unfiltered signal, and in the 1-50 Hz range in 5 Hz bins, averaged over channels. Features were normalized to their median value in the non-anesthetized portion of the recording, or lowest anesthetic concentration, if no artifact-free minute was available.

#### Regressors

Machine-learning analyses were performed using Python (Python Software Foundation, NH, USA) in the Spyder (Pierre Raybaut, The Spyder Development Team) development environment. Model training and performance evaluation were carried out using the scikit-learn toolbox. The set was iteratively (n=100) divided in a training (2/3 of the set) and a cross-validation (1/3) set. Hyper-parameter of the model were tuned on the training set, which was further split using the standard 3-fold cross-validation split implemented by the function “GridSearchCV”, to which a “pipeline” object was passed. The “pipeline” object was built using the “Pipeline” function, and concatenating quantile transformation of the input features (“Quantile Transformer”, tuning the number of quantiles), feature selection (“Select Percentile”, using mutual information and tuning the percentage of features to select) and Radial Basis Function (RBF) kernel support-vector classification/regression (tuning the regularization parameters C and epsilon (regression only), and the kernel coefficient gamma). The classifier input was fed to the regressor as an additional feature. Performance assessment was then computed on the cross-validation set. Regressor decision space was reduced and plotted with t-sne. The decision space was approximated by imposing a Voronoi tessellation on the 2d plot, using k-nearest regression on the t-sne coordinates (*57*).

#### Statistics

Statistical analyses were performed using R Statistical Software (Foundation for Statistical Computing, Austria). Data were tested for significant differences (*P<0.05, **P<0.01 and ***P<0.001) using non-parametric one- and two-way repeated-measures ANOVA (ARTool R package) with Bonferroni corrected post hoc analysis (emmeans R package). Correlations were computed using Spearman’s rank correlation coefficient (rho). No statistical measures were used to estimate sample size since effect size was unknown. For main experimental results, more information about tests used, values and parameters are provided in the supplementary material (Tab. S2).

## Acknowledgements

We thank P. Putthoff, A. Marquardt, A. Dahlmann, and K. Titze for excellent technical assistance. **Funding:** This work was funded by grants from the European Research Council (ERC-2015-CoG 681577 to I.L.H.-O.), the German Research Foundation (Ha 4466/10-1, SPP 1665, SFB 936 B5 to I.L.H.-O.) and the International Anesthesia Research Society (to L.C.). I. L. H.-O. is member of FENS Kavli Network of Excellence.

## Author contributions

M.C., S.H.B. and I.L.H.-O. designed the experiments, M.C., S.H.B., S.G., J.K.K., J.A.P., and L.C. carried out the experiments, M.C., S.H.B., S.G., J. K.K. and J.A.P. analyzed the data, M.C., S.H.B., L.C., C.B.B. and I.L.H.-O. interpreted the data and wrote the paper. All authors discussed and commented on the manuscript.

## Supplementary Materials

**Fig. S1.**
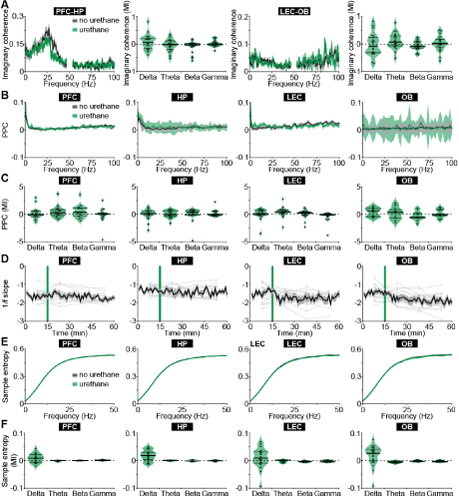
Urethane anesthesia does not affect spectral features and timing of activity in neonatal mice. (**A**) Line plots displaying the imaginary coherence between PFC-HP and LEC-OB in neonatal mice (P8-10) as a function of frequency before (black) and after (green) urethane injection. Violin plots displaying the MI of the imaginary coherence in delta (2-4 Hz), theta-alpha (4-12 Hz), beta (12-30 Hz) and gamma (30-100 Hz) frequency bands. (**B**) Line plots displaying the PPC of MUA to the oscillatory phase before and after urethane injection. (**C**) Violin plots displaying the MI of PPC in delta, theta, beta and gamma frequency bands. (**D**) Line plots displaying the slope of the 1/f decay for gamma frequencies over time. Green lines mark the time point of urethane injection. (**E**) Line plots displaying the sample entropy as a function of frequency before and after urethane injection. (**F**) Violin plots displaying the MI of the sample entropy in delta, theta, beta and gamma frequency bands.

**Fig. S2.**
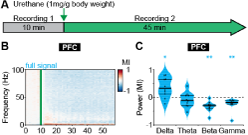
Frequency-specific effects of urethane anesthesia in juvenile mice. (**A**) Schematic representation of experimental paradigm of LFP recordings in PFC of non-anesthetized and urethane-anesthetized juvenile mice (P24-39). (**B**) Color-coded MI of oscillatory power for full signal before and after urethane injection. Green line corresponds to the time point of urethane injection. (**C**) Violin plots displaying the MI of oscillatory power in delta (2-4 Hz), theta-alpha (4-12 Hz), beta (12-30 Hz) and gamma (30-100 Hz) frequency bands for full signal.

**Fig. S3.**
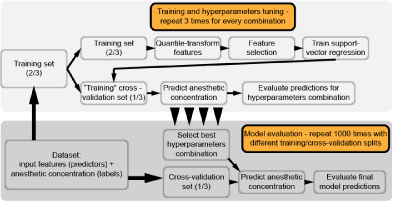
Machine learning algorithm. Flowchart depicting steps for machine learning algorithm.

**Fig. S4.**
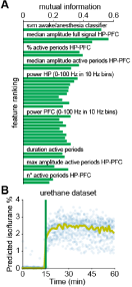
Median amplitude is most informative for predicting anesthetic concentration in neonatal mice. (**A**) Bar plot displaying the feature ranking for anesthesia depth prediction by mutual information between each feature and anesthesia depth. (**B**) Scatter plot displaying predicted isoflurane concentration using features of LFP recordings from PFC and HP of urethane-anesthetized mice. Green line marks the time point of urethane injection.

**Fig. S5.**
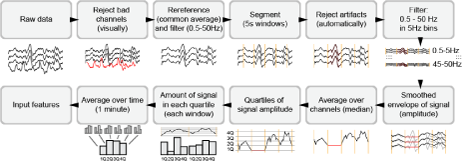
EEG data processing. Flowchart depicting analysis steps for EEG data processing.

**Fig. S6.**
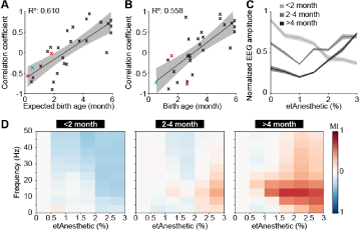
Age-dependent switch from broadband suppression to frequency-specific effects of general anesthesia on EEG activity for post conceptual age and frontal electrodes. (**A**) Scatter plot displaying the correlation coefficient of median EEG amplitude and anesthetic concentration in relationship to expected birth age for sevoflurane (black), isoflurane (red), and desflurane (blue). (**B**) Scatter plot displaying the correlation coefficient of median EEG amplitude of frontal electrodes and anesthetic concentration in relationship to birth age for sevoflurane (black), isoflurane (red), and desflurane (blue). (**C**) Line plots displaying normalized EEG amplitude of frontal electrodes as a function of anesthetic concentration. (**D**) Color-coded MI of median EEG amplitudes of frontal electrodes in different frequency bands as a function of anesthetic concentration for human babies of 0-2 months (left), 2-4 months (middle) and 4-6 months (right).

**Fig. S7.**
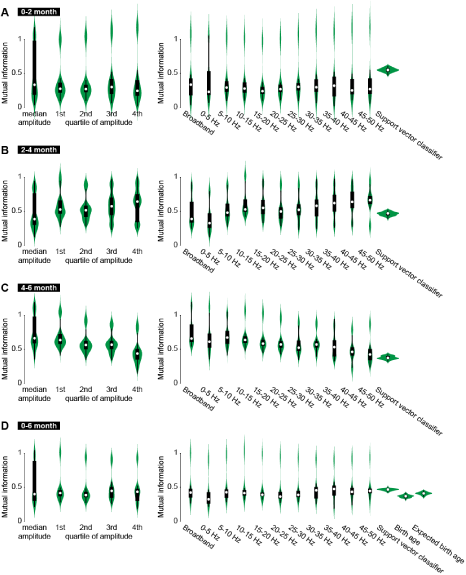
Features predicting anesthetic concentration from EEG recordings in human infants. (**A**) Violin plots displaying mutual information between each feature and predicted anesthetic concentration for amplitude-related features (left) and frequency-related features (right) for human infants of 0-2 months of age. (B) Same as (A) for human infants of 2-4 months of age. (C) Same as (A) for human infants of 4-6 months of age. (D) Same as (A) for human infants of 0-6 months of age.

**Tab. S1.** Demographic information.

|  | **All Subjects** | **Age Groups** |  |  |
| --- | --- | --- | --- | --- |
|  | **0-6 months** | **0-2 months** | **2-4 months** | **4-6 months** |
|  | **N=35** | **n=6** | **n=19** | **n=10** |
| Age at birth (weeks; median, IQR) | 38.00 [36.64, 39.00] | 39.71 [38.25, 40.53] | 37.00 [34.00, 39.00] | 39.00 [37.44, 39.00] |
| PNA (months; median, IQR) | 3.06 [2.64, 4.42] | 1.51 [1.36, 1.73] | 2.89 [2.74, 3.52] | 5.54 [5.08, 5.91] |
| Weight (kg, median, IQR) | 5.97 [4.89, 7.09] | 4.91 [4.80, 5.01] | 5.76 [4.89, 6.62] | 7.29 [6.42, 8.10] |
| Male (n, %) | 30 (85.7) | 5 (83.3) | 16 (84.2) | 9 (90.0) |
| Duration of Anesthesia (mins; median, IQR) | 114.00 [82.5, 181.00] | 190.00 [123.50, 211.50] | 114.00 [85.00, 154.00] | 89.00 [76.5, 181.75] |

| **ID** | **Age at birth (weeks)** | **Postnatal age (months)** | **Weight (kg)** | **Sex** | **Surgery** | **Duration Anesthesia** |
| --- | --- | --- | --- | --- | --- | --- |
| 1 | 39,0 | 0,53 | 3,7 | Female | Anorectoplasty | 217 |
| 2 | 38,0 | 1,35 | 5,7 | Male | Hernia Repair | 103 |
| 3 | 41,0 | 1,41 | 4,8 | Male | Hernia Repair | 93 |
| 4 | 40,4 | 1,61 | 5,0 | Male | Extrophy of Bladder Closure, Spica Cast | 185 |
| 5 | 40,6 | 1,77 | 4,8 | Male | Colostomy Closure | 268 |
| 6 | 37,5 | 1,87 | 5,0 | Male | Hernia Repair, Frenulotomy | 195 |
| 7 | 39,0 | 2,04 | 5,0 | Male | Circumcision | 60 |
| 8 | 42,0 | 2,60 | 6,0 | Male | Hernia Repair | 89 |
| 9 | 34,0 | 2,60 | 4,8 | Male | Hernia Repair | 103 |
| 10 | 29,1 | 2,69 | 3,3 | Male | Hernia Repair, Circumcision | 152 |
| 11 | 39,0 | 2,73 | 7,2 | Male | Hernia Repair, Orchidopexy | 187 |
| 12 | 30,3 | 2,76 | 3,4 | Male | Hernia Repair | 149 |
| 13 | 35,4 | 2,79 | 4,7 | Male | Hernia Repair, Meatoplasty | 114 |
| 14 | 34,0 | 2,83 | 5,0 | Male | Hernia Repair | 79 |
| 15 | 39,0 | 2,89 | 6,1 | Female | Hernia Repair | 96 |
| 16 | 37,0 | 2,89 | 6,3 | Male | Hernia Repair | 140 |
| 17 | 39,0 | 3,02 | 5,8 | Female | Vaginoscopy | 156 |
| 18 | 29,0 | 3,06 | 4,2 | Male | Hernia Repair | 185 |
| 19 | 38,0 | 3,45 | 6,9 | Male | Hernia Repair | 72 |
| 20 | 39,0 | 3,52 | 7,2 | Male | Fistulotomy | 58 |
| 21 | 37,0 | 3,52 | 6,1 | Female | Nephrectomy | 170 |
| 22 | 37,0 | 3,58 | 5,7 | Male | Hernia Repair | 118 |
| 23 | 41,0 | 3,71 | 5,8 | Male | Hernia Repair | 102 |
| 24 | 39,0 | 3,81 | 7,0 | Male | Pyleoplasty | 177 |
| 25 | 39,0 | 3,81 | 7,6 | Male | Fistulotomy | 81 |
| 26 | 39,0 | 4,04 | 7,4 | Male | Hernia Repair | 94 |
| 27 | 42,0 | 4,80 | 6,3 | Female | Hernia Repair | 76 |
| 28 | 29,1 | 5,03 | 7,8 | Male | Hernia Repair | 78 |
| 29 | 38,0 | 5,26 | 6,3 | Male | Hypospadias Repair | 22 |
| 30 | 39,0 | 5,36 | 8,5 | Male | Orchidopexy | 76 |
| 31 | 39,1 | 5,72 | 7,2 | Male | Colostomy Closure | 310 |
| 32 | 29,0 | 5,78 | 9,1 | Male | Fistulotomy | 62 |
| 33 | 40,4 | 5,95 | 8,2 | Male | Hypospadias Repair | 189 |
| 34 | 39,0 | 6,01 | 6,7 | Male | Chordee Release | 160 |
| 35 | 30,3 | 6,05 | 3,2 | Male | Circumcision | 84 |

**Tab. S2.**
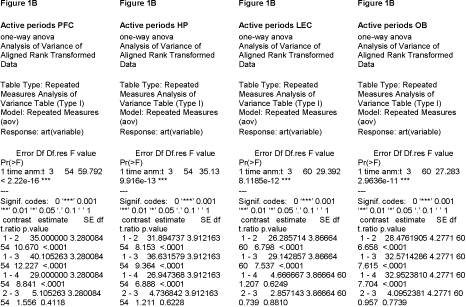
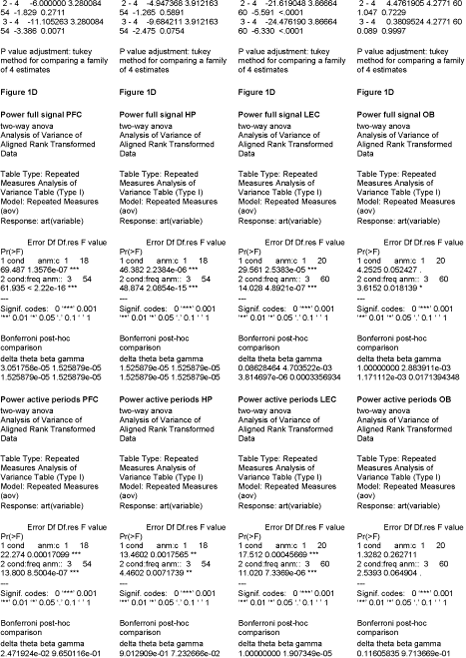
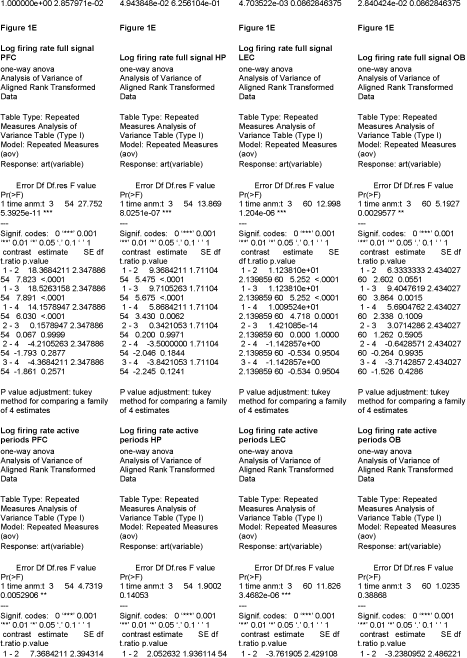
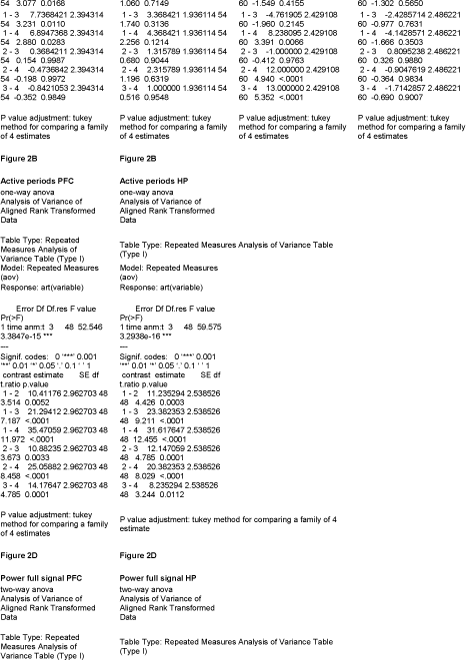
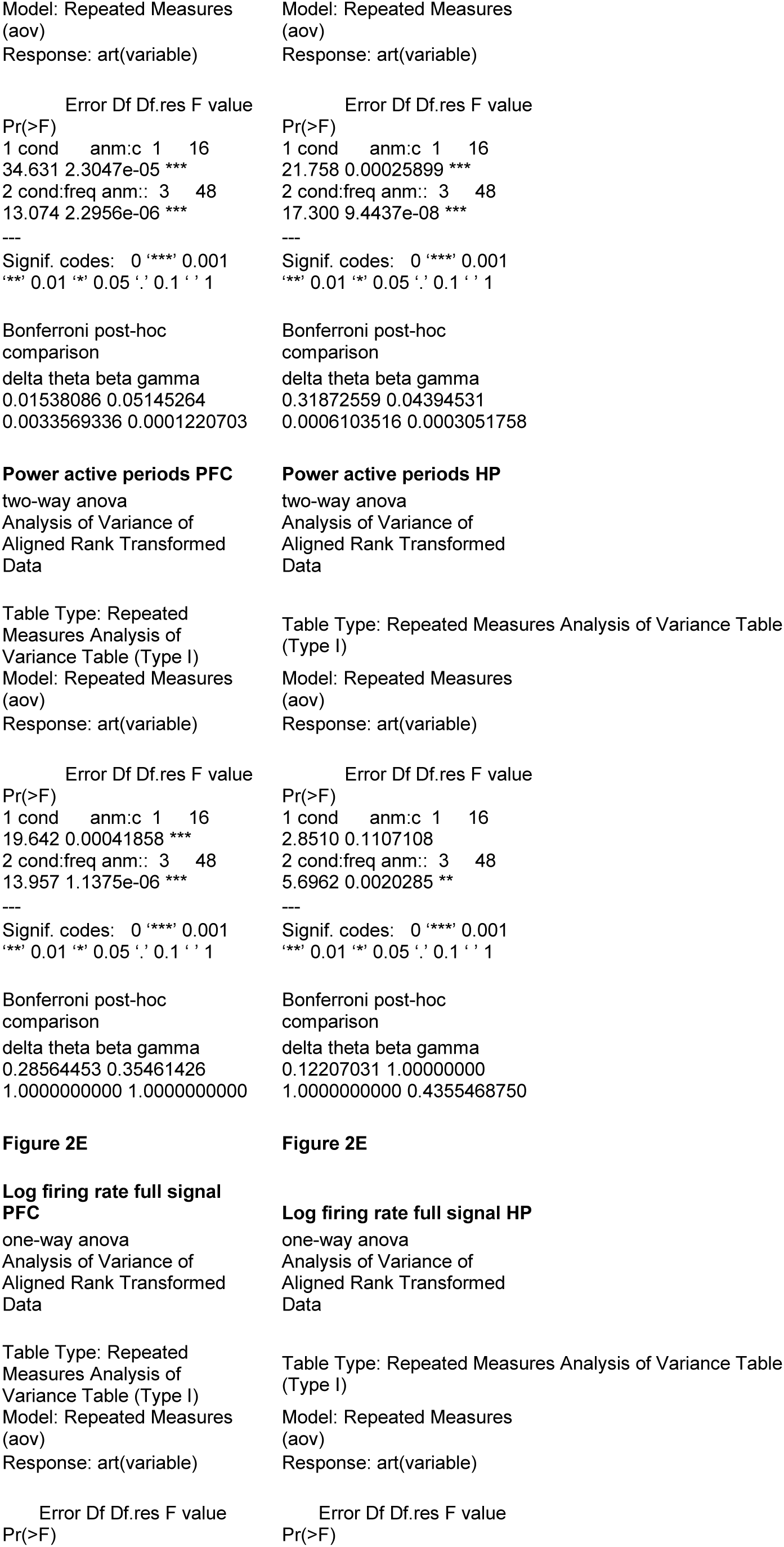
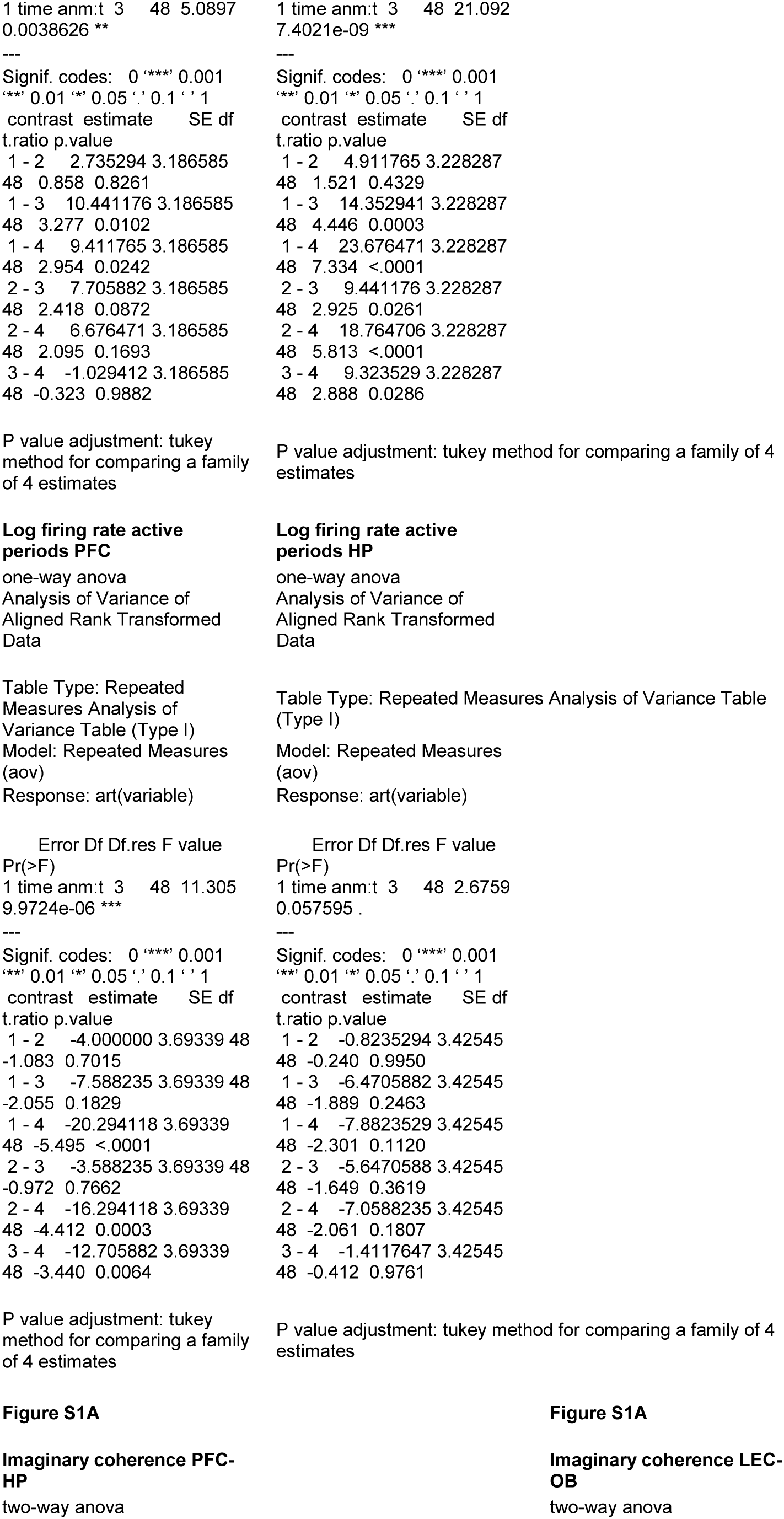
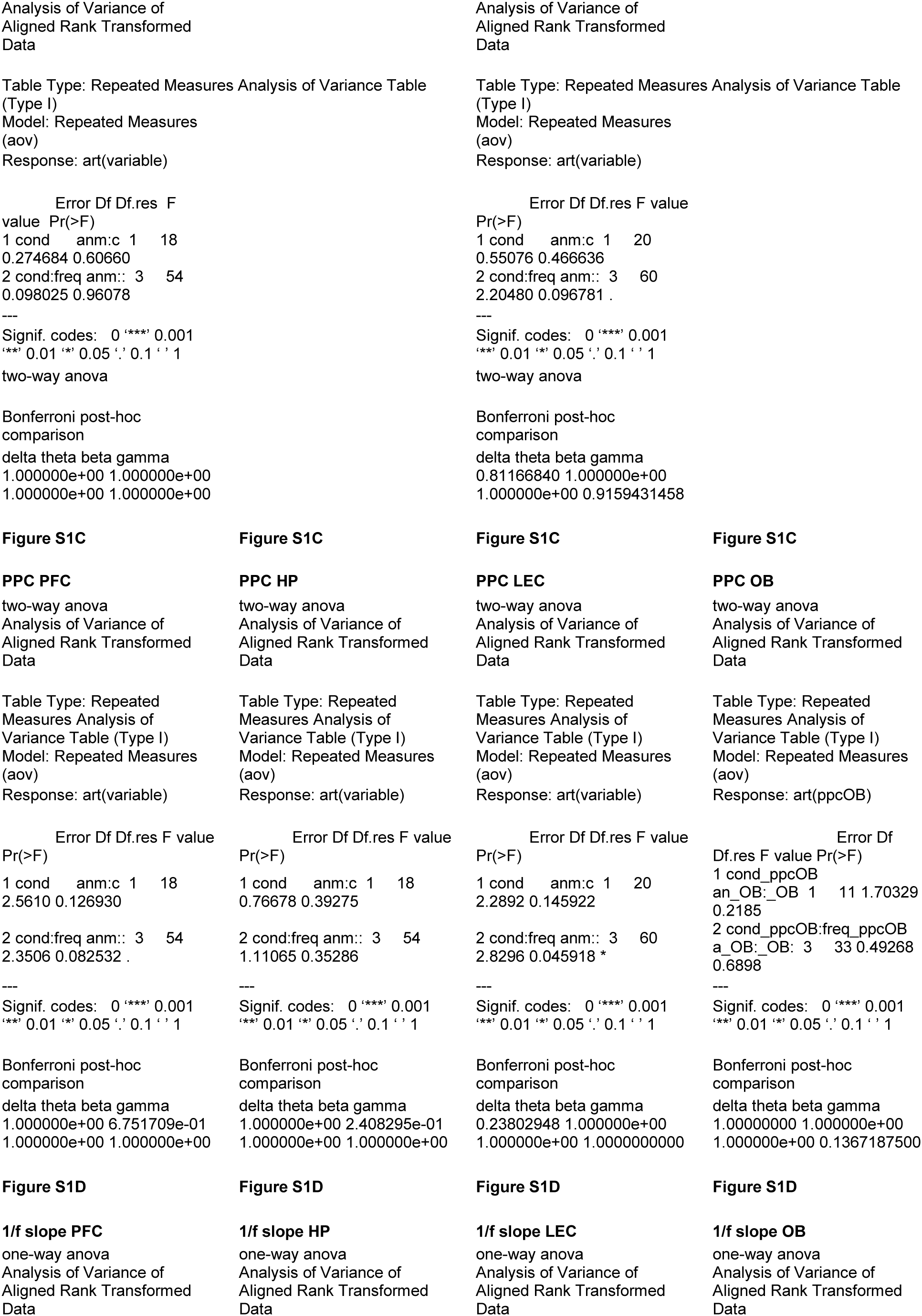
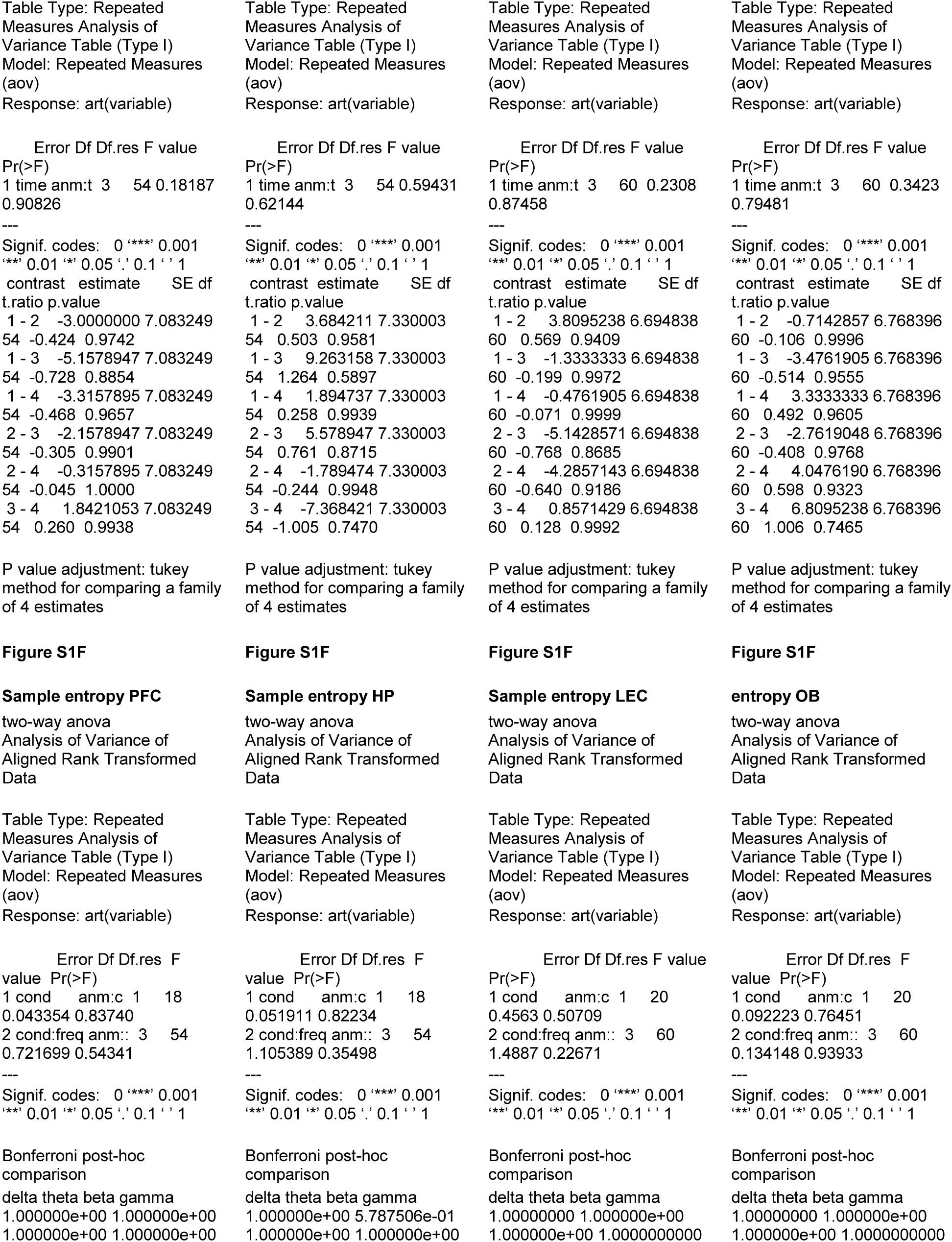
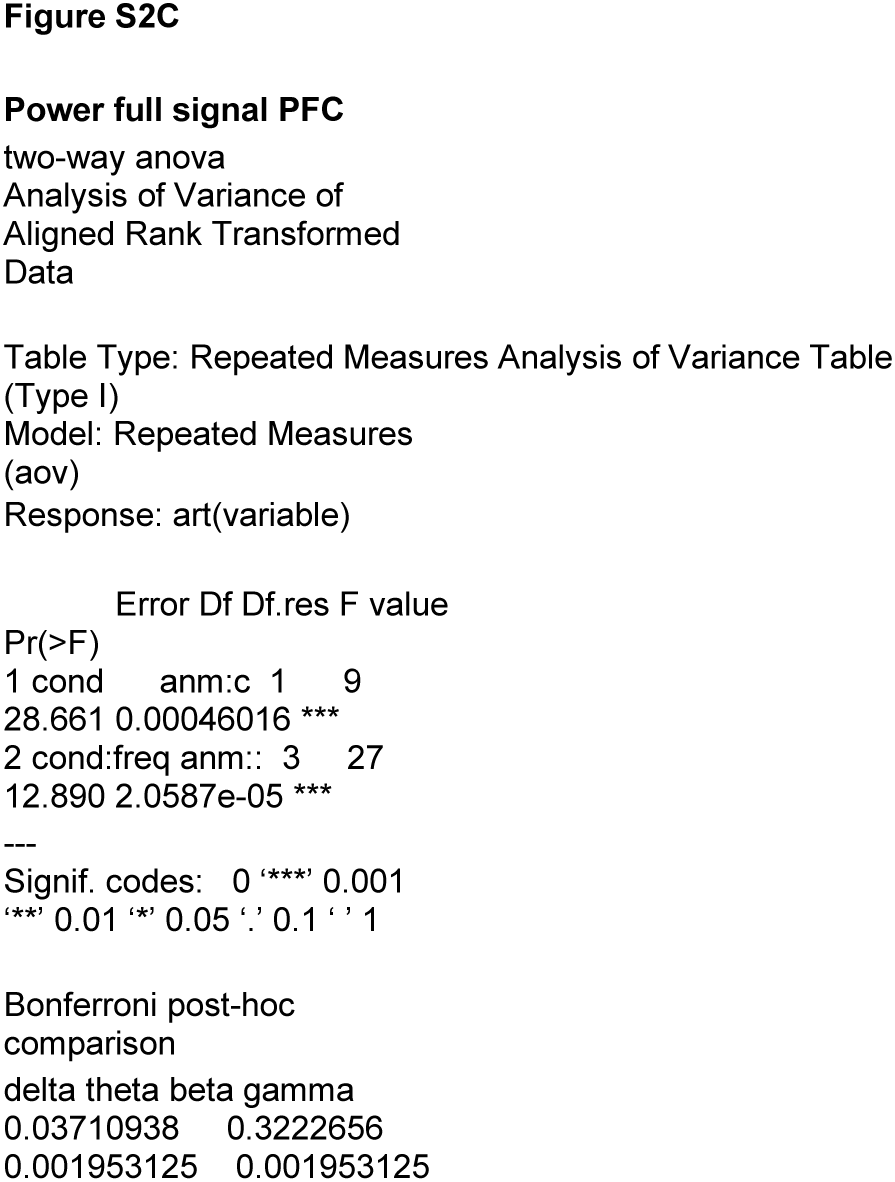
Statistics summary.

